# *tmap*: topological analysis of population-scale microbiome data

**DOI:** 10.1101/396960

**Authors:** Tianhua Liao, Yuchen Wei, Mingjing Luo, Guoping Zhao, Haokui Zhou

## Abstract

Population-scale microbiome study poses specific challenges in data analysis, from enterotype analysis, identification of driver species, to microbiome-wide association of host covariates. Application of advanced data mining techniques to high-dimensional complex dataset is expected to meet the rapid advancement in large scale and integrative microbiome research. Here, we present *tmap*, a topological data analysis framework for population-scale microbiome study. This framework can capture complex shape of large scale microbiome data into a compressive network representation. We also develop network-based statistical analysis for driver species identification and microbiome-wide association analysis. *tmap* can be used for exploring variations in a population-scale microbiome landscape to study host-microbiome association.

**Availability and implementation:** *tmap* is available at GitHub (https://github.com/GPZ-Bioinfo/tmap), accompanied with *online documentation* and tutorial (http://tmap.readthedocs.io).

**Contact:** http://hk.zhou@siat.ac.cn

## 1 Introduction

Population-scale microbiome studies have led to the identification of host factors associated with gut microbiome (Falony et al., 2016), characterization of gut microbiome enterotypes and their driver microbial species (Arumugam et al., 2011). The application of microbiome-wide association analysis (**MWAS**) to population cohorts has revealed marker species associated with diseases (Gilbert et al., 2016). Standard microbiome data analysis relies on ordination techniques and regression analysis to discover variations of microbiome community and identify their association with host phenotypes (Knight et al., 2018). With the trends in conducting population-scale studies, multi-omics data integration and meta-analysis, rapidly advances in microbiome analysis methods and standards are expected, and will benefit from machine learning techniques in analyzing large scale high dimensional dataset (Mallick et al., 2017).

Topological data analysis (**TDA**) provides a promising technique for analyzing large scale complex data. The most popular *Mapper* algorithm is effective in distilling data-shape from high dimensional space, and provides a compressive network representation (Singh *et al.*, 2007; Lum *et al.*, 2013). This algorithm has demonstrated its superior to traditional dimension reduction methods in analyzing biological and medical data, in which many of the challenges are similar to microbiome studies, such as subtyping of disease (Li et al., 2015), clustering of cancer groups, and identification of associated gene features (Nicolau *et al.*, 2011).

Here, we present the *tmap* software as an implementation of the **TDA** *Mapper* framework for population-scale microbiome data analysis. We developed *tmap* to enable easy adoption of TDA in microbiome data analysis pipeline, providing network-based statistical methods for enterotype analysis, driver species identification, and microbiome-wide association of host meta-data. As demonstrated in this study, by re-analyzing the FGFP microbiome data (Falony et al., 2016), *tmap* is a promising microbiome data analysis framework for both large scale exploratory analysis and network-based hypothesis testing in population-scale microbiome study.

## 2 Workflow and implementation

### 2.1 *tmap* workflow

We implemented *tmap* as a Python package, consisting of modules and classes for major steps of the *Mapper* algorithm (Fig.1). Design of the modules and classes was motivated to provide flexible and consistent application programming interfaces (**API**s) for the workflow (online documentation). By using the APIs, the workflow can be extended to incorporate machine learning methods from other Python packages, such as *Scikit-learn* (Pedregosa et al., 2011). Furthermore, *tmap* APIs allow for easy integration of the workflow into other microbiome analysis pipelines, such as QIIME (Caporaso *et al.*, 2010). *tmap* has several advantages in microbiome data analysis, from the use of precomputed beta-diversity distance matrix, helper functions for TDA parameter selection, to the downstream network-based species enrichment and meta-data association analysis (online documentation). *tmap* provides an integrated and streamlined workflow to take the results from standard microbiome data analysis pipelines (such as QIIME) to advanced analysis and generates insights from microbiome data.

**Fig. 1.**
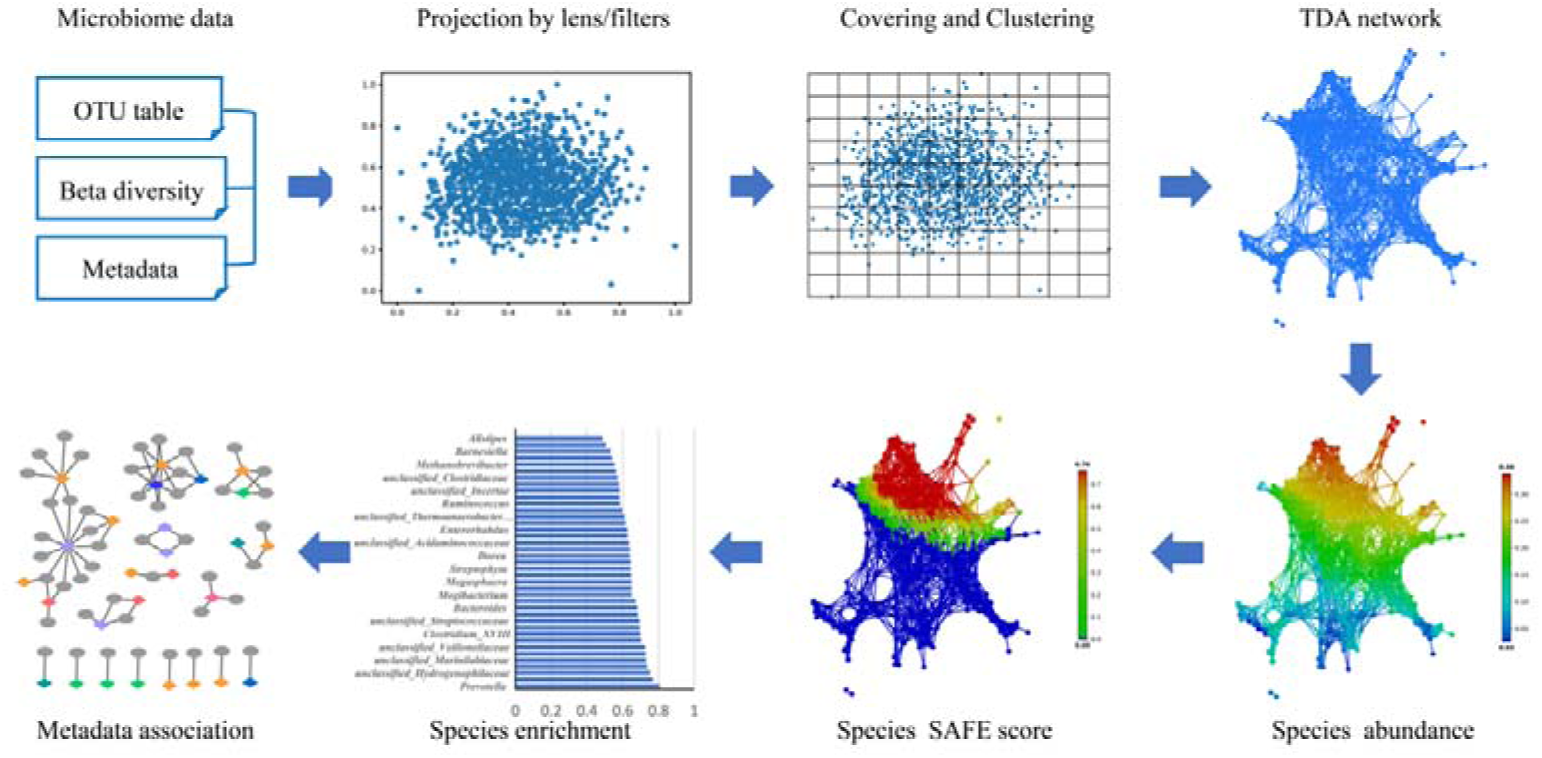
*tmap* workflow and steps of topological analysis of microbiome data. Starting with inputs of microbiome data (OTU table, beta-diversity distance matrix and sample metadata), *tmap* proceeds with data projection, covering, clustering and TDA network construction. Identification of driver species and association analysis of metadata are based on the TDA network structure, by calculating the SAFE network enrichment score.

### 2.2 Steps of *tmap* analysis

A typical *tmap* analysis consists of four major steps, as illustrated in Fig.1. The first step is to project microbiome data into a low-dimensional space using dimension reduction methods with a specified distance metric. The second step is to make topological covering and clustering of the projected data. Depending on sample size and required resolution of microbiome variations, parameters of covering and clustering should be chosen and examined carefully. We provide a detailed explanation and guideline in the online documentation for these parameters. The third step is to generate, visualize and explore the microbiome TDA network. This step allows for discovering of patterns in microbiome variations and for visualizing how the pattern changes in the network along with meta-data, by mapping colors on the network. At last, network analysis is performed to identify driver species responsible for the observed community variation, and for metadata association analysis.

### 2.3 Network enrichment analysis of driver species

Enterotypes and driver species can be identified from microbiome data using ordination and regression analysis. In *tmap*, we used a network enrichment technique for driver species analysis, based on a network representation of microbiome variations, which is adapted from the spatial analysis of functional enrichment (SAFE) algorithm (Baryshnikova, 2016) (Supplementary Material). Our approach successfully recovered all the top ten driver species from the FGFP cohort. Our approach also identified new driver species that are separated into multiple node clusters, which are also enriched in the network (Supplementary Table 2 and Fig. S1). The multi-cluster pattern of these driver species could be hard to detect with linear regression methods and therefore missed from the FGFP study. Another advantage of network enrichment analysis is a direct assignment and visualization of enterotype for each node (Supplementary Fig. S2).

### 2.4 Network-based microbiome-wide association analysis

*tmap* provides a novel alternative to standard MWAS by using network-based association analysis. Instead of association with individual samples, *tmap* uses SAFE enrichment scores at node level, which is a group of samples of highly similar microbiome composition. Application of this approach to the FGFP cohort successfully identified most of the reported associations, with improved power and effect sizes (Supplementary Table 4). We have also identified new associations not reported in the original study (Supplementary Fig. S4).

## 3 Conclusion

*tmap* is an integrated and advanced framework for topological analysis of microbiome data for population-scale study. The framework captures the topological shape of microbiome composition and variation into a network representaion, followed by network-based statistical analysis. We expect that *tmap* will enable researchers to obtain more insights from microbiome data, from enterotype analysis, driver species identification, understanding of microbiome landscape, to microbiome-wide association analysis.

## Acknowledgements

We thank Huansheng Gao and Peijian Qiu for their help in improving the Python codes for network plotting and coloring.

